# Antagonistic Pleiotropy Is Unexpectedly Rare In New Mutations

**DOI:** 10.1101/301754

**Authors:** Mrudula Sane, Joshua John Miranda, Deepa Agashe

## Abstract

Pleiotropic effects of mutations may underlie diverse biological phenomena such as ageing and specialization. In particular, antagonistic pleiotropy (“AP”: when a mutation has opposite fitness effects in different environments) generates tradeoffs, which may constrain adaptation. Models of adaptation typically assume that AP is common – especially among large-effect mutations – and that pleiotropic effect sizes are positively correlated. The rare empirical tests of these assumptions have largely focused on beneficial mutations observed under strong selection, whereas most mutations are actually deleterious or neutral, and are removed by selection. We quantified the incidence, nature and effect size of pleiotropy for carbon utilization across 80 single mutations in *Escherichia coli* that arose under mutation accumulation (i.e. weak selection). Although ~46% of the mutations were pleiotropic, only 11% showed AP, which is lower than expected given the distributions of fitness effects for each resource. In some environments, AP was more common in large-effect mutations (but not synergistic pleiotropy, SP); whereas pleiotropic effect sizes were positively correlated for SP (but not AP). Thus, AP is generally rare, is not consistently enriched in large-effect mutations, and often involves weakly deleterious antagonistic effects. Our unbiased quantification of mutational effects therefore suggests that antagonistic pleiotropy is unlikely to cause maladaptive tradeoffs.

## INTRODUCTION

Biologists have long observed that organisms maximize resource allocation to one trait while compromising allocation to another trait (Goethe)(Lenoir 1984). Such tradeoffs manifest as negative correlations between traits, and may constrain evolution by limiting the breadth of phenotypes available to organisms (Rees 1993). The nature and strength of tradeoffs between traits can thus dictate whether organisms evolve to be generalists or specialists (Ferenci 2016). Tradeoffs also underlie diverse biological phenomena such as life-history strategies (Zera and Harshman 2001; Sgrò and Hoffmann 2004), ageing (Kirkwood 2005), and assembly of microbial communities and host-microbe interactions (Litchman et al. 2015). Although tradeoffs in resource allocation are undeniable, they remain relatively poorly understood at the mechanistic level. Tradeoffs can occur when multiple neutral or deleterious mutations accumulate and degrade traits under weak selection, leading to a negative correlation with other traits evolving under positive selection (Elena and Lenski 2003). For instance, in Lenski’s long term experimental evolution lines, bacteria evolving under strong selection for one metabolic function (growth on glucose) lost multiple other metabolic functions because selection on these traits was very weak, allowing deleterious mutations to accumulate (Cooper 2014; Leiby and Marx 2014). Alternatively, tradeoffs may occur when a single mutation increases fitness in a specific environment (or trait), simultaneously reducing fitness in alternate environments (or a second trait) (Cooper and Lenski 2000). Such mutations are antagonistically pleiotropic for the two traits or environments, and the phenomenon is called antagonistic pleiotropy (henceforth “AP”).

The evolutionary impact of AP clearly depends on its incidence and magnitude. If AP is frequent or involves large-effect mutations, the resulting tradeoffs are more likely to constrain adaptation. Historically, models of adaptive evolution have assumed that AP is the predominant form of pleiotropy (Otto 2004, Lande 1983), implying that synergistic pleiotropy (SP; when a mutation simultaneously either increases or decreases fitness in two different environments) is relatively uncommon. However, for single beneficial mutations in *Escherichia coli*, AP between fitness on glucose and alternate carbon sources was rare compared to positive SP (Ostrowski et al. 2005). Similarly, most of the first-step beneficial mutations isolated from laboratory-evolved *E. coli* populations showed SP, while only a few were strongly antagonistically pleiotropic (Dillon et al. 2016). Thus, contrary to model assumptions, empirical data suggest that AP may not be the predominant form of pleiotropy. A second assumption of theoretical models is that large effect mutations are more predisposed to show AP (Lande 1983), potentially explaining the prevalence of small effect mutations during adaptation in natural populations (Lande 1983; Orr 1992; Dillon et al. 2016). However, no empirical study has explicitly tested this assumption. Finally, the pleiotropic effect size of mutations is assumed to be proportional to their fitness effect in the selective environment where the mutation arose, i.e. its primary effect size (Orr and Coyne 1992). Contrary to this assumption, previous studies found that the antagonistic effect size was not correlated with the primary effect size (Ostrowski et al. 2005; Dillon et al. 2016). Taken together, empirical studies indicate that SP is more common than AP, at least among beneficial mutations. Additionally, the direct and pleiotropic effects of beneficial mutations appear to be positively correlated when pleiotropy is synergistic, but not when pleiotropy is antagonistic. Thus, widely used models of adaptive evolution make assumptions that are either empirically untested or are poorly supported. Although the empirical studies mentioned above provided important results, all of them focused on beneficial mutations, which represent only a small fraction of all mutations. The majority of mutations are expected to be either neutral or mildly deleterious (Eyre-Walker and Keightley 2007; Bataillon and Bailey 2014). Thus, by focusing only on beneficial mutations, we ignore most of the distribution of fitness effects of mutations (henceforth “DFE”), in turn ignoring the role of non-beneficial mutations in evolution. During adaptive evolution – depending upon the strength of selection and population size – neutral and mildly deleterious mutations may accumulate and later influence adaptation to new environments. For example, deleterious alleles can persist in a population at low frequencies as standing genetic variation, but rise to high frequencies in new favourable environments (Steiner et al. 2007; Barrett and Schluter 2008).

To obtain unbiased estimates of AP, we evolved replicate populations of *E. coli* under mutation accumulation (henceforth “MA”) for multiple generations on a rich medium (Fig. 1). This regime of experimental evolution minimizes the strength of selection due to repeated bottlenecking of the populations, allowing all but lethal mutations to accumulate. We sequenced several time points frozen during experimental evolution to identify lines that had a single mutation relative to their immediate ancestor. Across 38 MA lines, we identified 80 isolates carrying new single mutations (including single nucleotide changes and small indels < 10 bp; henceforth “mutants”) relative to their immediate ancestor. To determine the incidence of AP (i.e. the proportion of mutants that showed increased fitness on resource A and decreased fitness on resource B), we measured the growth rate of each of these mutants and their respective mutational ancestors on 11 different carbon sources. For each pair of resources, we compared the observed incidence of AP with null distributions generated by randomly sampling from the independent DFEs for each resource (Fig 1). We find that while pleiotropy is not rare among new mutations, AP is quite uncommon and variable across resources, even when compared to the null distribution. Although the incidence of AP often increases with the effect size of the mutation, the form of the relationship varies across resources. Finally, we find that the fitness effect sizes of mutations showing AP are either uncorrelated or negatively correlated. Taken together, our results suggest that AP is more rare than previously thought, indicating that AP-mediated tradeoffs are generally unlikely to constrain adaptation.

**Figure 1:**
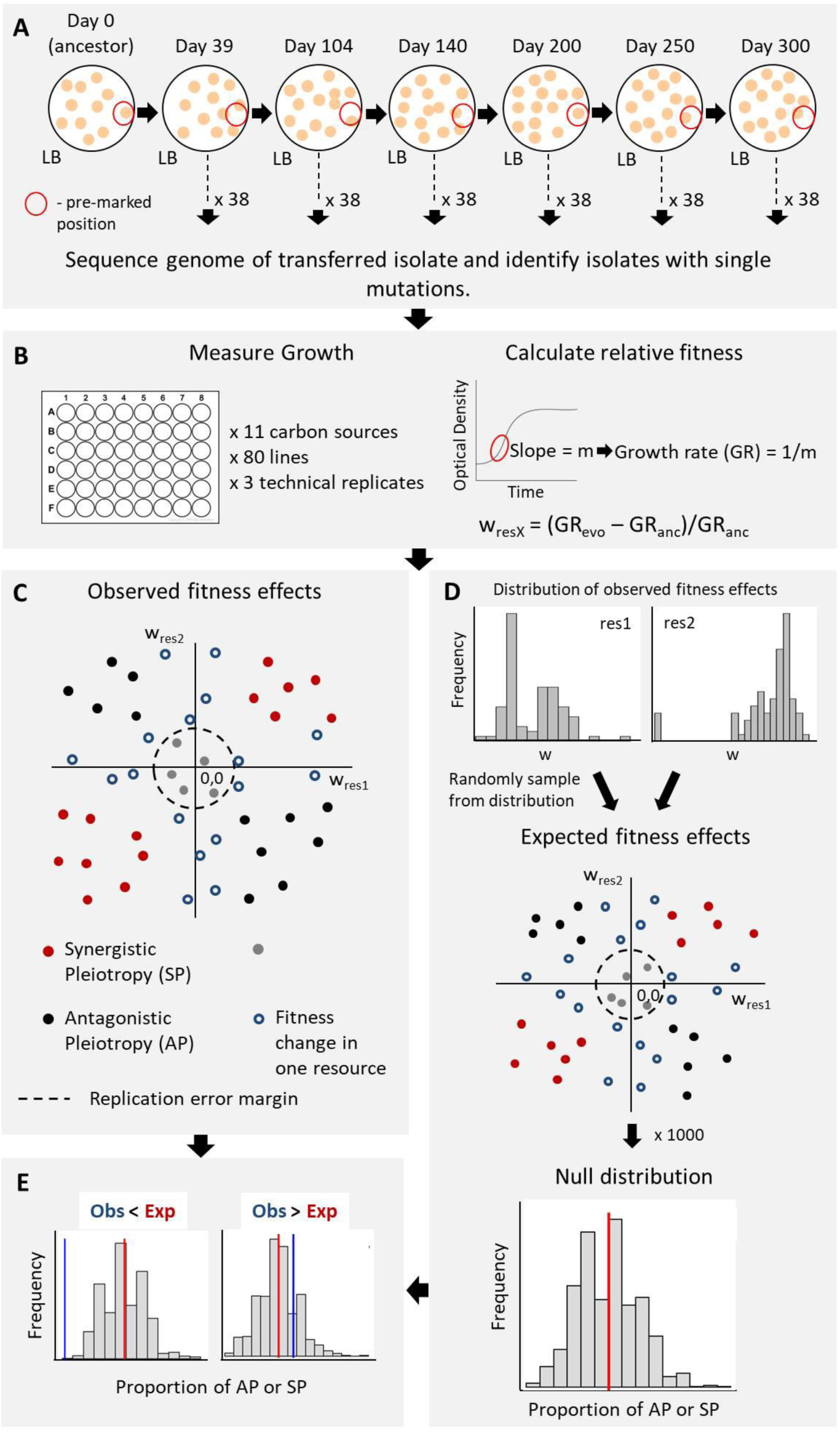
Experimental design and analysis. (A) Mutation accumulation experiment (B) Estimating fitness effects of single step mutations in different environments (C) Calculating the observed incidence of pleiotropic mutations (D) Generating null distributions for the incidence of pleiotropic mutations, given independent distributions of fitness effects in each environment (E) Comparing the difference in observed vs. expected incidence of pleiotropy for each resource pair.

## MATERIALS AND METHODS

### Bacterial strains and media

We obtained the wild-type strain of *Escherichia coli* K-12 MG1655 from the Coli Genetic Stock Centre (CGSC, Yale University). We streaked this strain on an LB agar plate, picked one colony at random, made glycerol stocks, and used this isolate as the wild-type (WT) in all subsequent experiments. For fitness assays, we used liquid culture media: LB broth (Miller, Difco), or M9 minimal salts medium + 5mM of a carbon source (glucose, trehalose, fructose, maltose, lactose, galactose, succinate, pyruvate, melibiose, malate, fumarate; Sigma-Aldrich).

### Mutation Accumulation (MA) lines

We suspended the ancestral WT colony in 200μL of LB broth and streaked out 2μL of the suspension on LB agar plates (with 2% agar), founding 38 replicate lines with two lines per Petri plate. We incubated plates at 37°C for 24 hours, and propagated the MA lines by picking a random colony from every line every 24 hours and streaking it onto a fresh agar plate. We ensured random picking of colonies by choosing a well-isolated colony closest to a pre-marked spot on the plate for each line. Every 4-5 days, we made frozen stocks of each line by inoculating a part of the transferred colony in LB broth at 37°C for 2-3 hours and freezing 1mL of the growing culture with 8% DMSO at −80°C. For the current study, we used time-points frozen on days 39, 104, 140, 200, 250 and 300. The protocol is summarized in Fig. 1A.

### Whole genome sequencing to identify single-step mutations

We inoculated 2μL of each frozen stock (and ancestors of each strain) in 2mL LB and allowed the cells to grow overnight at 37°C with shaking at 200rpm. We extracted genomic DNA using the GenElute Bacterial Genomic DNA kit (Sigma-Aldrich), quantified genomic DNA using the Qubit HS dsDNA assay (Invitrogen), and prepared genomic DNA libraries for each isolate using the Nextera XT DNA Library Preparation Kit (Illumina), using the manufacturer’s instructions in each case. We sequenced libraries on the Illumina Hi-seq 2500 platform using either 2×100bp paired-end reaction chemistry (giving an average coverage of 150× per genome; range 102× to 240×) or the 1×100 single-end reaction chemistry (giving an average coverage of 70× per genome; range 21× to 140×). We discarded reads with quality scores less than Q30, retaining >95% reads per genome. We aligned filtered reads to the NCBI reference *Escherichia coli* K-12 MG1655 genome (RefSeq accession ID GCA_000005845.2) using the Burrows-Wheeler short-read alignment tool BWA (Li and Durbin 2009). We used SAMtools to process the BWA output and generate pileup files (Li et al. 2009), and used the VARSCAN package to extract the list of SNPs and short indels (<10bp) (Koboldt et al. 2009). To identify long indels (> 10bp length), we used Breseq (Deatherage and Barrick 2014). We discarded all mutations that occurred at <80% frequency or were supported by <10 reads on both strands. Compared to the NCBI reference genome, our WT ancestor contained one SNP (single nucleotide change) and one indel that were present in all evolved MA lines, and were hence discarded from further analysis. We did not find any long indels in our sequenced isolates. The mutations observed in the MA lines will be described in detail in a separate publication.

For this study, we identified all isolates carrying a single mutation with respect to their immediate ancestor. For instance, if a line had one mutation on day 39 and an additional mutation at day 200, we retained both these isolates for further analysis, but discarded intermediate time points (days 104 and 140) since they did not represent single mutational steps. In this case, we obtained two distinct single-mutation steps from the same line: for the evolved isolate at day 39, we considered the WT as the ancestor; and for the evolved isolate at day 200, we considered the evolved isolate at day 39 as the ancestor. If a line already had two mutations on day 39 (the first frozen time point) and four mutations in the subsequent time point (day 104), we discarded all these isolates from further analysis since we could not find single-mutation steps for this line. Details of the 80 isolates representing single mutational steps (“mutants”) are given in Table S1.

### Measuring growth rate as a fitness measure

We measured the fitness of all mutants, along with their respective ancestors (see summary in Fig. 1B). We inoculated each mutant from its freezer stock into M9 minimal salts medium with 0.4% glucose, kept at 37°C with shaking at 200 rpm for 16 hours. We then inoculated 6μL of this culture into 594μL growth media (LB broth or M9 minimal salts medium + 5mM carbon source) in 48-well plates (Costar), incubated in a shaking tower (Liconic) at 37°C. Plates were read by an automated growth measurement system (Tecan, Austria) every 40 minutes for 18 hours. We tested the growth rate of three technical replicates per line per carbon source; meaningful biological replicates could not be obtained since we had a single colony at the end of each MA line transfer. In each 48 well plate, we included the WT and ancestor of the MA lines being tested as controls, and we used these to estimate variation in growth rates across plates and to calculate relative fitness of evolved isolates. We estimated maximum absolute growth rates using the Curve Fitter software (Delaney et al. 2013). For a subset of 40 mutants, we repeated fitness assays in glucose, galactose, and pyruvate to ensure that growth rates were consistent across independent runs (Fig. S1).

### Estimating fitness effect sizes and calculating observed and null estimates of AP and SP

For each mutant,we calculated relative fitness as: (Growth rate of mutant – Growth rate of ancestor)/Growth rate of ancestor (see Fig. 1B), using the average for three technical replicates. A negative value indicated that growth rate had decreased compared to the ancestor, while a positive value indicated increased growth rate compared to the ancestor. Growth rates for WT measured in different plates run on different days varied by less than 5%. Similarly, the error in measurement of growth rates across technical replicates (run on the same day) was also less than 5%. To account for this variability, we considered mutants with <5% change in fitness from the ancestor as showing no change. For each pair of carbon sources, we calculated the proportion of mutants showing evidence of AP (relative growth rate <-0.05 in carbon source A but relative growth rate >0.05 in carbon source B) or SP (relative growth rate <-0.05 in both carbon source A and carbon source B as synergistic decreases in fitness; relative growth rate >0.05 in carbon source A and B as synergistic increases in fitness) (Fig 1C). To determine the proportion of comparisons showing AP or SP for each focal resource (Fig. 2), we calculated the total number of mutants showing AP or SP across all pairwise combinations with the focal resource. Because there were 10 possible resource pairs for each focal resource and 80 mutants for which AP or SP was measured, there were a total of 800 comparisons (10 resource pairs × 80 lines) for each isolate. Thus, we calculated the “observed” proportion of comparisons showing AP or SP for each focal resource as the number of mutants showing AP or SP, divided by 800.

**Figure 2:**
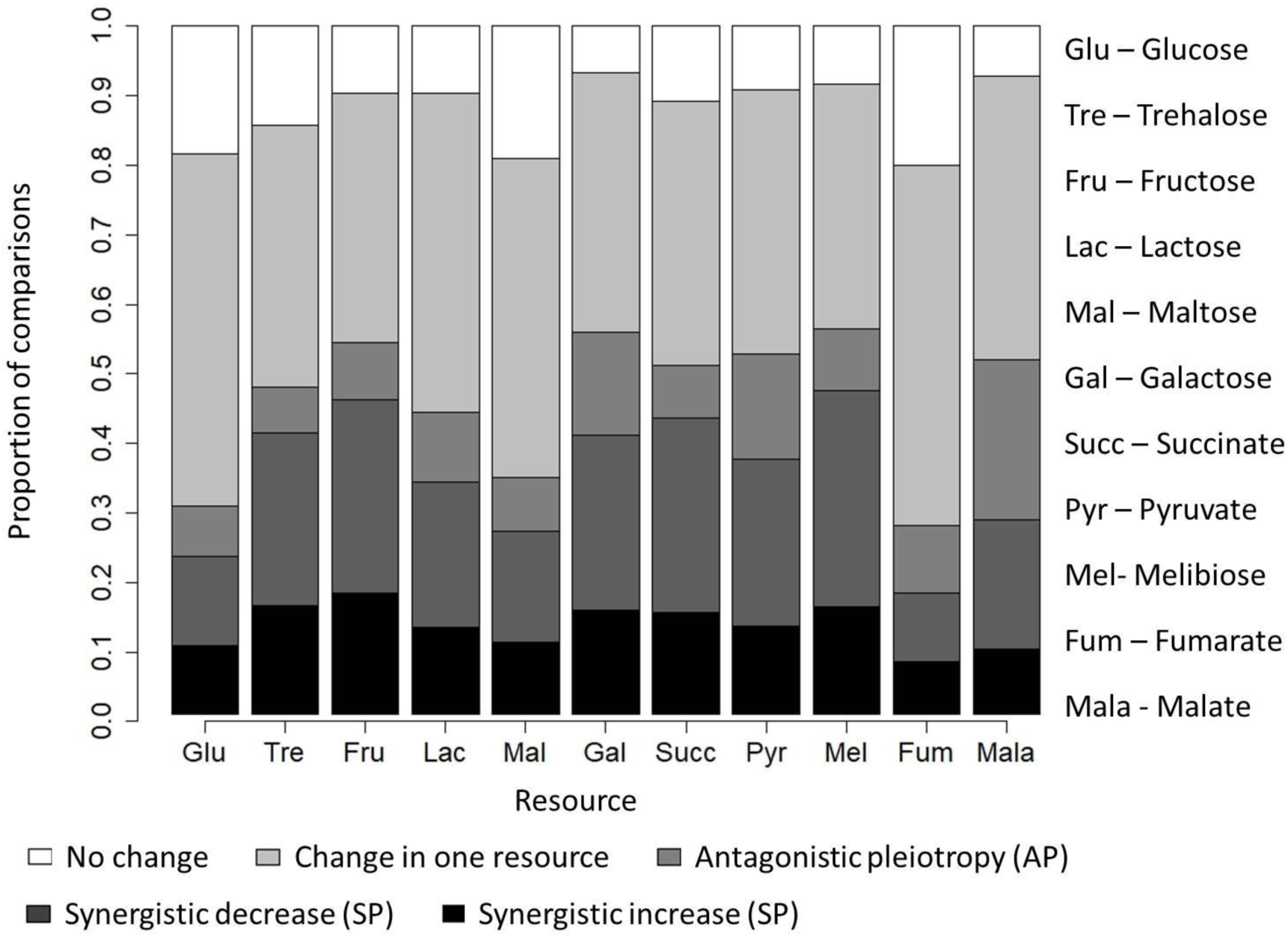
Incidence of pleiotropy among single mutational steps in wild type *E. coli.* Stacked bar plots show the observed proportion of mutants showing pleiotropy, change in fitness in only one resource, or no fitness change. Each bar represents pooled data across all pairwise resource comparisons for the focal resource indicated on the x-axis (80 mutants × 10 resource combinations involving the focal resource = 800 data points per bar).

To generate null distributions of the incidence of pleiotropy among all mutations for each of the 55 possible resource combinations, we randomly picked a fitness value from the observed distribution of fitness effects (DFE) for one resource, simultaneously picking a fitness value from the DFE for the other resource. We picked 80 such pairs of fitness values, and calculated the proportion of pairs showing AP or SP. We performed 1000 iterations of this process to generate a null distribution of the incidence of AP or SP for each resource pair. When generating null distributions of proportions of pleiotropy for beneficial mutations, for each resource pair (A and B), it was possible for a beneficial mutations to occur in either resource A or resource B. We accordingly generated two separate null distributions for each resource pair, leading to a total of 110 null distributions (Fig. 1D). For each null distribution, we estimated the average proportion of AP (or SP) as the “expected” incidence of AP (or SP), for comparison with the observed incidence of AP (or SP) for the specific resource pair (Fig. 1E).

### Testing for a correlation between the incidence of pleiotropy and fitness effect size

For isolates showing AP or SP, we used absolute values of relative fitness as a measure of the magnitude of pleiotropic effect size. We calculated the correlation between fitness effect size and proportion of pleiotropy in two ways. (1) We categorized absolute fitness effects (i.e. without considering the direction of fitness change) for each focal resource into four arbitrary classes: very low (relative fitness 0.05 – 0.1), low (relative fitness 0.1 – 0.2), medium (relative fitness 0.2 – 0.3), and high (relative fitness 0.3 – 0.4). We then counted the number of instances of pleiotropy (AP or SP, as appropriate) in each class. We then asked if the proportion of pleiotropy was correlated with the fitness class. (2) We selected only those mutants that showed pleiotropy (AP or SP) for each focal resource in turn. We then classified them into the four fitness effect bins based on their pleiotropic effect size, and counted the number of mutants falling in each class. Using these data, we asked: conditional to the occurrence of pleiotropy, how is it distributed across the four fitness effect classes? Similarly, to calculate the null expectation for the relationship between fitness effect size and proportion of pleiotropy, we binned, as described above, fitness values randomly drawn from the DFEs for individual resources. We measured the proportion of pleiotropy (AP or SP) within the null distribution, and asked if it was correlated with the fitness effects for each of the 55 resource pairs.

### Testing for a correlation between primary and pleiotropic fitness effect sizes

For each resource pair, we computed the Spearman’s rank correlation between the magnitudes of effect sizes (absolute values of relative fitness, as above) in the two resources, for all mutants that showed pleiotropy (AP or SP, as appropriate). We included fitness data for LB in this analysis, as LB is the environment in which our MA lines evolved. Thus, for this analysis, the number of resources is 12, and the number of resource pairs is 66. We excluded those resource pairs from analysis in which less than 5 mutants showed the specific type of pleiotropy. Since AP is rare, we could therefore compute effect-size correlations for 50 of 66 resource pairs. For SP, we computed effect-size correlations for all 66 resource pairs.

## RESULTS

### Antagonistic pleiotropy is rare, and varies across environments

To estimate the incidence of pleiotropy, we measured the fitness effect (growth rate) of single mutations obtained during an MA experiment, on 11 different carbon sources (Fig. 1). As expected, the distribution of fitness effects (DFEs) observed for each resource showed that on average, ~49% of all sampled mutations were deleterious, and would have been missed if we focused only on beneficial mutations (Fig. S2). Combining data across all mutants and resource pairs (80 mutants x 55 resource pairs = 4400 data points), we observed pleiotropy in ~46% of the cases (Fig. 2). However, most pleiotropic mutations were synergistic (SP, ~35% of total) rather than antagonistic (AP, ~11%). Importantly, resource identity had a significant impact on the incidence of both AP and SP (Fig. 2; p < 0.05, generalized linear model with binomial errors; Tables S2 and S3; also see Tables S4 and S5 for all pairwise resource comparisons). Malate had the highest incidence of AP (~23%), while melibiose showed the highest incidence of SP (50%). Overall, AP was relatively rare compared to SP.

Another way to quantify the incidence of pleiotropy is to ask whether a given mutation shows pleiotropy across multiple resource pairs. Most mutations (72 of 80) showed AP for at least one pair of resources, with a median of 6 and a range of 0-24 resource pairs (out of 55 total resource pairs; Fig S3). In contrast, all mutants showed SP for at least one resource pair, with a median of 16 resource pairs (Fig. S3). These results again highlight the relative rarity of AP compared to SP. The relatively high frequency of SP suggests that the paucity of AP cannot be explained by a general inability to simultaneously detect small, pleiotropic fitness effects in multiple environments.

Finally, we compared the observed incidence of AP and SP with the null expectation derived from DFEs for each resource in a given resource pair combination (Fig. 1C-E). Using random, repeated sampling from observed DFEs for each resource pair, we estimated that the expected incidence of AP was ~16 % (average across all resource pairs; Fig. S6); this is greater than the observed incidence of ~11% described above. For each resource pair, we tested whether the observed proportion of mutants showing AP was significantly greater or lower than expected from the null distribution for the specific resource pair. We found that for most resource pairs (39 of 55), significantly fewer mutations showed AP than expected by chance (Table 1; Fig S6). In contrast, in most cases SP was observed significantly more often than expected (46 of 55 resource pairs; Table 1; Fig. S7). When we considered only beneficial mutations for each focal resource, the pattern for AP was even more stark, with all 110 resource pairs showing lower AP incidence (on average, ~3.9% across all resource pairs) than expected (average ~40% across all resource pairs) (Table 1; see also Fig. S8). However, for beneficial mutations, SP showed a reverse pattern than for all mutations, with 109 of 110 resource pairs showing less SP (~13% across all resource pairs) than expected (~26% across all resource pairs) (Table 1; see also Fig. S9). Together, these results reinforce our conclusion that AP is very rare in new mutations. In contrast, SP is more common than expected, except when considering only beneficial mutations. Overall, our results may explain why AP-mediated tradeoffs have been difficult to uncover in empirical studies: AP is not only rare, but also depends on the environment.

**Table 1:**
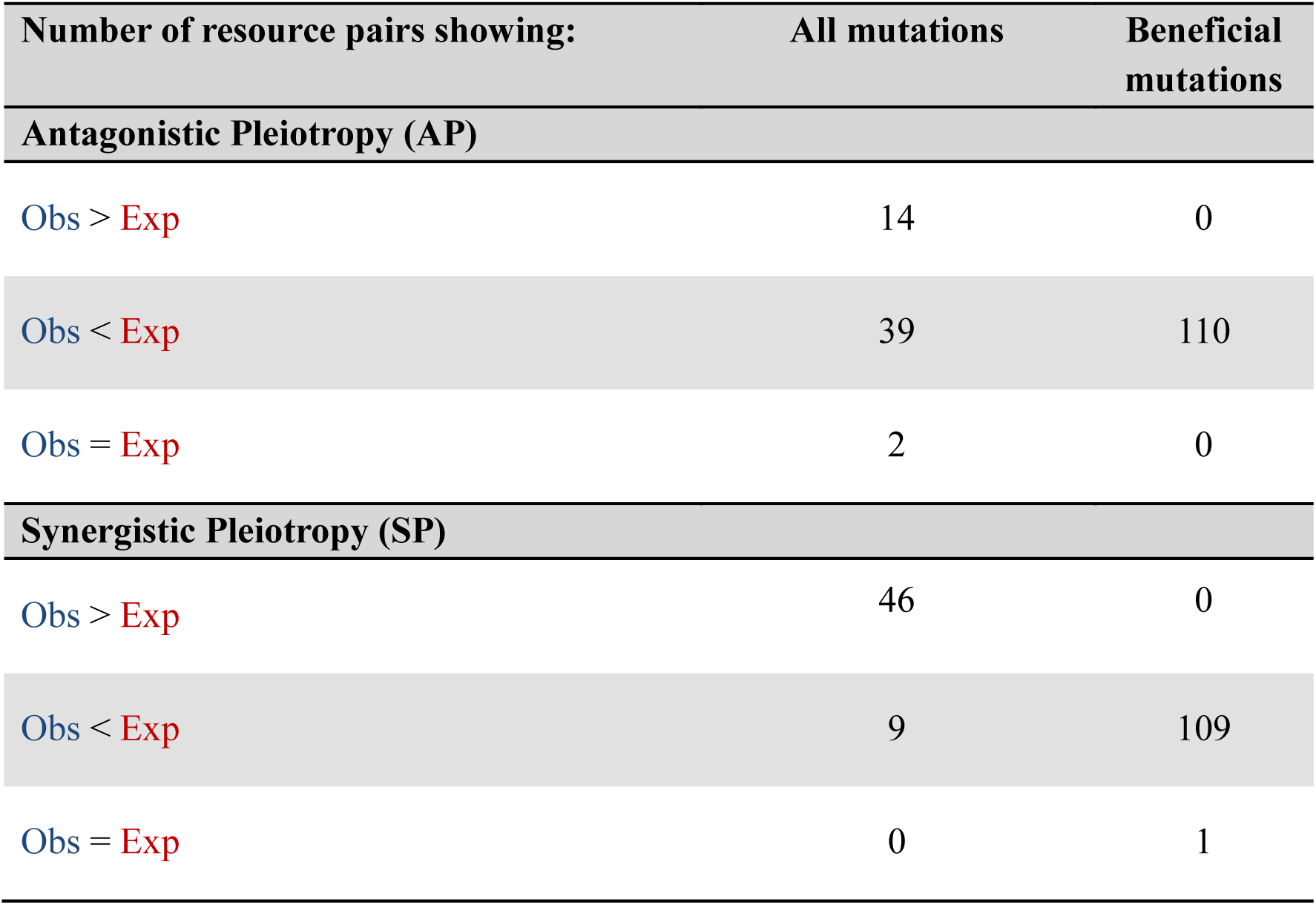
Summary of the number of resource pairs that show specific patterns of observed vs. expected incidence of pleiotropy for all mutations, or only for beneficial mutations. Observed and expected proportions of AP or SP were designated as significantly different (Obs > Exp or Obs < Exp) when p < 0.05 for a Student’s t-test comparing the observed proportion of AP or SP with the mean of the null distribution for each resource pair (see Fig. 1C-E). Null distributions of the incidence of AP and SP for each resource pair, and observed incidences, are shown in Fig. S7 and Fig. S8.

| Number of resource pairs showing: | All mutations | Beneficial mutations |
| --- | --- | --- |
| **Antagonistic Pleiotropy (AP)** |  |  |
| $\text{Obs} > \text{Exp}$ | 14 | 0 |
| $\text{Obs} < \text{Exp}$ | 39 | 110 |
| $\text{Obs} = \text{Exp}$ | 2 | 0 |
| **Synergistic Pleiotropy (SP)** |  |  |
| $\text{Obs} > \text{Exp}$ | 46 | 0 |
| $\text{Obs} < \text{Exp}$ | 9 | 109 |
| $\text{Obs} = \text{Exp}$ | 0 | 1 |

### Large-effect mutations are more likely to show pleiotropy

Theoretical models of adaptation assume that large-effect mutations are more commonly associated with pleiotropic effects, and that these pleiotropic effects are mostly deleterious (Lande 1983). To test this hypothesis, for each focal resource we grouped fitness effect sizes into four arbitrary classes: very low (relative fitness 0.05 – 0.1), low (relative fitness 0.1 – 0.2), medium (relative fitness 0.2 – 0.3), and high (relative fitness 0.3 – 0.4). Across all resources, ~37%, 45%, 14% and 4% of fitness effects were classified in the respective classes. We then tested the relationship between the incidence of AP and fitness effect size in two ways.

We first asked: in each of the four fitness effect size classes, what proportion of mutants show AP? Considering each focal resource in turn, we observed distinct relationships between the proportion of AP and the mutational effect size. Four resources showed the predicted, monotonic positive correlation (Kendall’s rank correlation, p < 0.05; first row in Fig. 3A; Table S6); three resources showed a concave positive relationship (second row in Fig. 3A); lactose showed a significant negative correlation; and the remaining three resources did not show a significant correlation between the incidence of AP and the fitness effect size. The correlation patterns for 7 of 11 resources supported the prediction that large-effect mutations are more likely to show AP; but the form of this relationship was not consistent across resources. Since a large fraction of mutations (37%) fall within the smallest effect size class, the relatively low incidence of AP in this bin is consistent with the conclusion that AP is generally rare. For SP, we observed more consistent relationships: the incidence of SP was positively correlated with effect size class for 10 of 11 focal resources (Fig. S5A).

**Figure 3:**
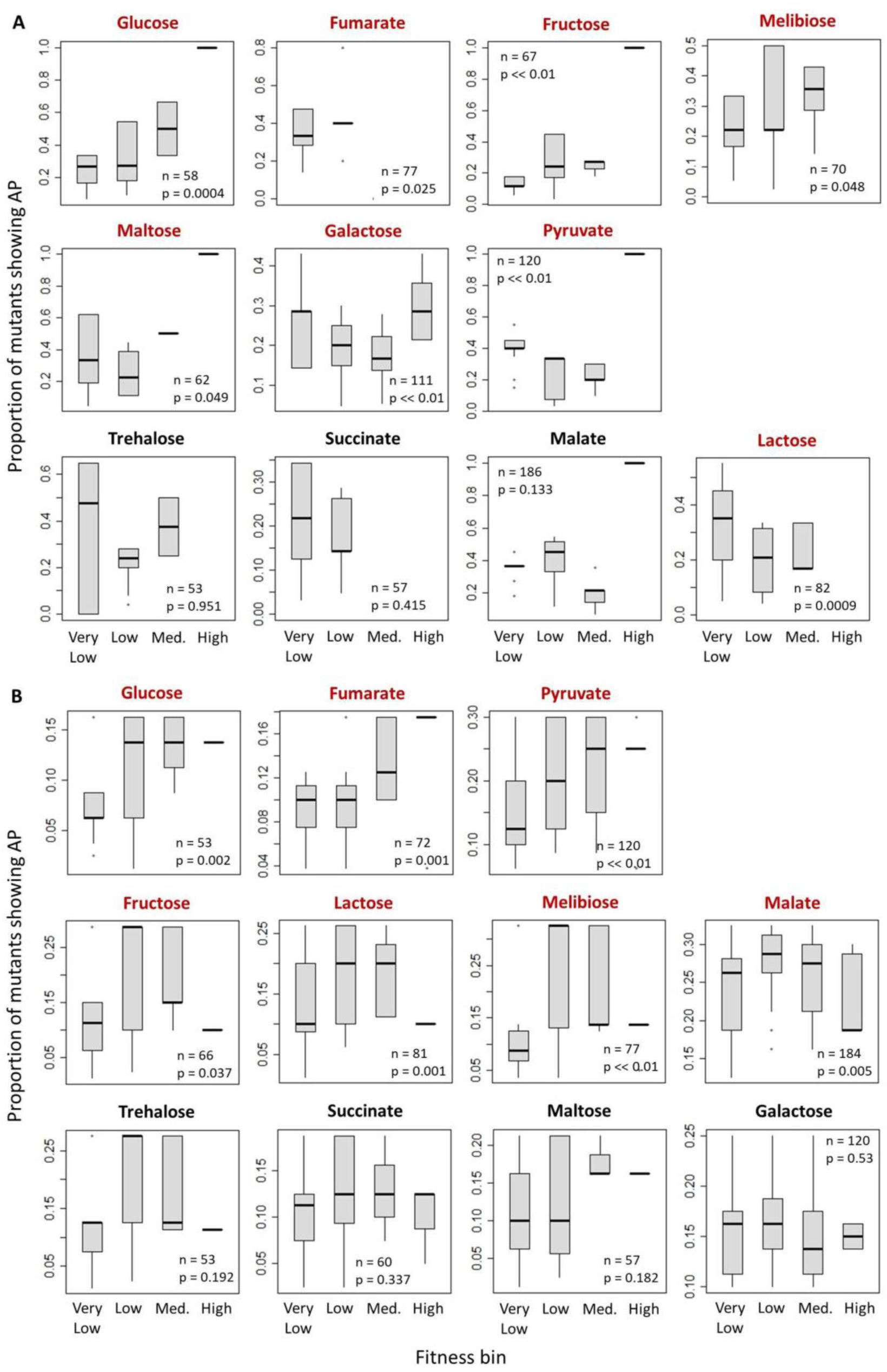
Relationship between the incidence of AP and fitness effect size. (A) Proportion of mutants showing AP in each fitness effect size class. For each mutant, we classified measured fitness effects for the focal resource into effect size classes, and then calculated the proportion of measurements representing AP with any other resource. Each plot thus represents a total of 80 measurements, of which *n* (indicated in each plot) represented AP. (B) Distribution of AP-causing mutations across fitness effect size classes. We filtered all instances of AP across our dataset (total 943), of which *n* (indicated in each plot) represented data for a given focal resource. For each resource, we calculated the proportion of measurements belonging to each fitness effect size class. Plot titles in red indicate a significant correlation between the fitness effect size class and the incidence of AP (p <0.05, Kendall’s rank correlation; see inset p values in each panel; also see Tables S6 and S8). Corresponding results for the correlation between expected AP incidence (based on null distributions) and fitness effect size are shown in Fig. S10.

Next, we asked: conditional on the occurrence of AP, do antagonistically pleiotropic mutations occur more frequently in large effect size classes? We again found variable patterns across resources: three resources showed a monotonic or saturating increase (first row, Fig 3B); four resources showed a convex relationship with highest AP incidence at intermediate fitness effect sizes (second row, Fig. 3B); and the remaining four resources showed no correlation (Table S8). In contrast, for datasets generated from randomly sampling DFEs for each resource, we found that effect sizes were consistently negatively correlated with the proportion of AP (Fig S10; Table S9). Thus, the observed positive relationship between proportion of AP and effect size cannot be explained by a greater chance of detecting AP in large-effect mutations. A similar analysis for SP showed that 4 of 11 resources showed a positive correlation between effect size and incidence of SP (Fig. S5B; Table S10), compared to the null expectation of a consistently negative correlation (Fig. S11, Table S11). Thus, while the incidence of AP in observed mutations is often positively correlated with the fitness effect size of those mutations, this pattern is not generally true for SP.

Together, these results offer partial support for the prediction that large-effect mutations may be more like to show AP, with the caveat that the results vary dramatically across environments. For AP involving glucose, we observed a consistent, strong positive correlation in both analyses (compare Figs 3A and 3B), indicating that AP-mediated tradeoffs for glucose are more likely to occur for large-effect mutations. However, for other resources, the relationship between effect size and AP incidence is either inconsistent, insignificant, or more complex with intermediate maxima or minima. Hence, with respect to model assumption, this relationship is not robust and requires more careful attention.

### Primary fitness effects sizes are correlated with synergistic, but not antagonistic effect sizes

We tested the relationship between primary and pleiotropic effect sizes for our set of random mutations, measuring primary effect sizes in LB, the growth medium in which our MA lines evolved. We measured secondary effect sizes in M9 minimal medium + 5 mM single carbon sources as above. Contrary to expectation, we found that for AP, in most cases the primary fitness effect sizes (in LB) were uncorrelated with the secondary effect sizes in specific carbon sources (bottom row, Fig. 4A; Table S12). Thus, the magnitude of fitness change in LB is unrelated to fitness change in other resources. For pairwise comparisons across single carbon sources, all significant correlations (25 of 39 possible comparisons; ~64%) were negative (Fig. 4A). Thus, a large benefit in one carbon source was often associated with a small deleterious effect in another carbon source, or vice versa. Overall, antagonistic pleiotropic mutations either do not exhibit correlated fitness effects, or show negatively correlated fitness effects in different environments. Synergistic pleiotropic effect sizes were also uncorrelated with primary effect sizes in LB (Fig. 4B; Table S13), suggesting that changes in fitness in a rich medium such as LB may generally not be related to fitness on individual carbon sources. However, all other pairwise resource combinations were strongly positive (Fig. 4B), indicating that large-effect beneficial (or deleterious) mutations in one carbon source also had a large benefit (or disadvantage) in another carbon source. Thus, the predicted positive effect size correlations hold for synergistic, but not antagonistic pleiotropic effects.

**Figure 4:**
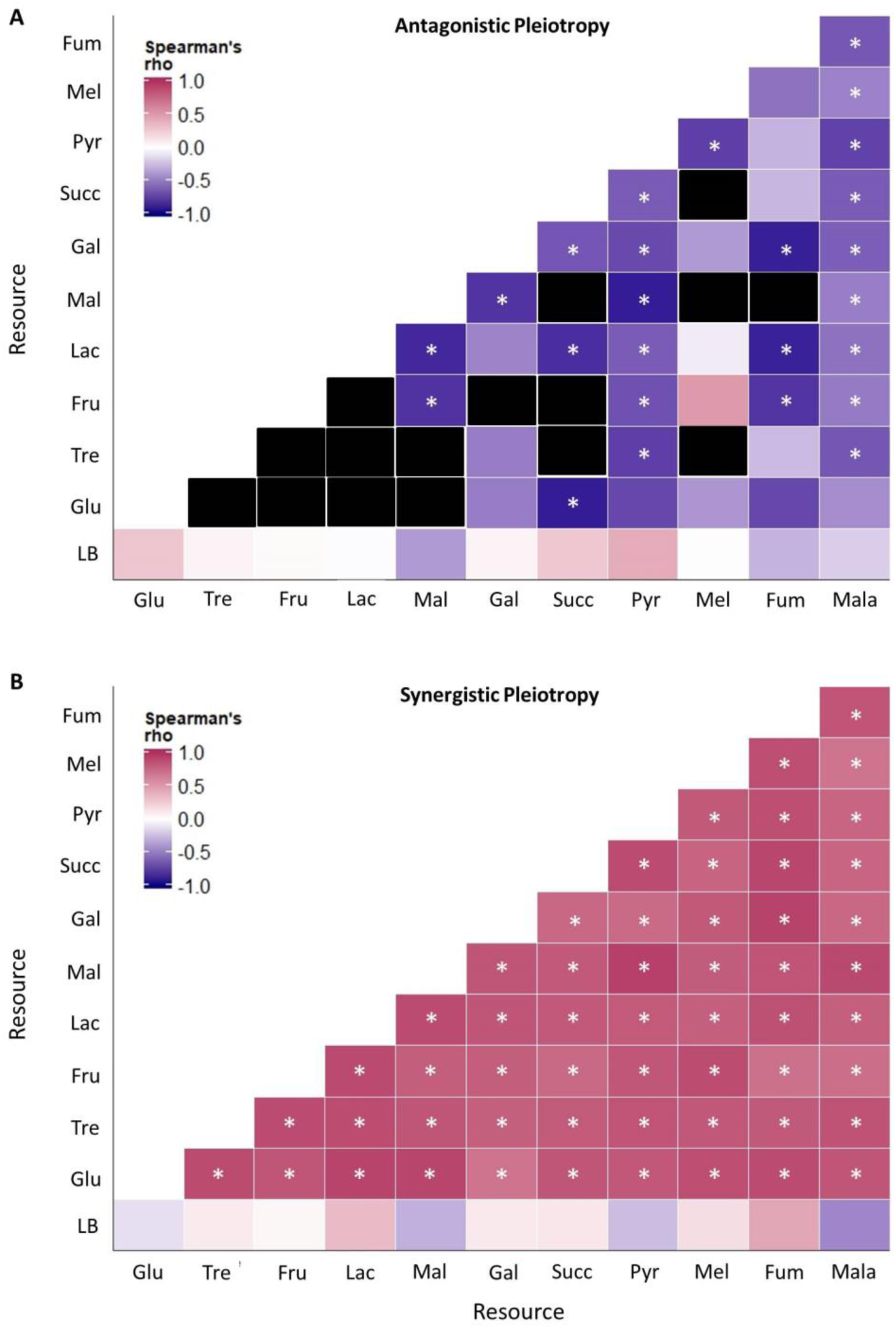
Correlation between primary and pleiotropic mutational effect size for each focal resource. Colored blocks indicate the Spearman’s rank correlation coefficient between a given focal resource (x-axis) vs. all other resources (y-axis), for mutants that showed (A) AP or (B) SP between a given pair of resources. In panel A, black blocks represent cases where correlations could not be computed because very few isolates showed AP (<5). Asterisks indicate a significant correlation between the magnitude of primary and pleiotropic effects (p < 0.05). Sample size, Spearman’s rho, and p values for each pairwise resource combination are given in Tables S12 and S13.

## DISCUSSION

In his artificial breeding experiments, Darwin observed Goethe’s Law of Compensation in action, stating “if nourishment flows to one part or organ in excess, it rarely flows, at least in excess, to another part” (Darwin 1859). This concept of tradeoffs has played a central role in evolutionary thinking. Tradeoffs influence most major ecological and evolutionary processes (Agrawal et al. 2010), including speciation and adaptive radiation (Kneitel and Chase 2004), evolution of specialization (Bono et al. 2017; Elena 2017), evolution of life histories (Stearns 1977, 1989), and assembly and coexistence in ecological communities (Tilman 2000; Bohannan et al. 2002). In bacteria alone, tradeoffs affect many key physiological processes (reviewed in Ferenci 2016): nutrient utilization and metabolism, antibiotic resistance (see also Hershberg 2017), resistance to phages, resistance to environmental stress, virulence and genome maintenance. However, the mechanisms underlying such phenotypic tradeoffs remain relatively poorly understood (Stearns 2000). A key mechanism is antagonistically pleiotropic mutations that can generate tradeoffs (Elena and Lenski 2003), but experimental measurements of the incidence, nature and effect size of pleiotropic mutations are rare. Here, we present a systematic analysis of pleiotropic fitness effects of a large, unbiased sample of single mutations observed in *E. coli* populations evolving under weak selection.

Our results provide three clear lines of evidence suggesting that AP due to single mutations is unlikely to be an important mechanism generating tradeoffs that hinder adaptation. First, we find that AP is generally rare in new mutations. In fact, among beneficial mutations, AP is much rarer than expected, indicating that beneficial mutations fixed during adaptation are unlikely to reduce fitness in other environments. Previous studies also found that only 10-14% of ~20 beneficial mutations showed AP (Ostrowski et al. 2005; Dillon et al. 2016). Second, we find that large-effect mutations are more likely to show AP in some (but not all) environments. Hence, AP may impose a major constraint only in specific environments and when adaptation involves large-effect mutations. Finally, we find that antagonistically pleiotropic mutations often have negatively correlated fitness effects, such that a highly beneficial mutation in one environment is only weakly deleterious in an alternate environment. Thus, such mutations are unlikely to impose a significant fitness disadvantage in new habitats. Together, our results contradict the prevalent idea that tradeoffs generated by AP may often constrain adaptation.

Our analysis of 80 randomly sampled single mutational steps has several advantages over previous studies. First, we determined the expected distribution of the proportion of AP given the underlying distributions of fitness effects in different carbon sources, providing a general framework to determine the occurrence of AP by chance alone. This null distribution allowed us to determine that the observed proportion of AP is significantly lower than the expected proportion of AP for ~71% of all resource pairs. A second advantage of our experiment is that we measured fitness effects in 11 distinct carbon sources (55 resource pairs), which is a much larger set of environments than previous analyses. This allowed us to detect many more instances of pleiotropy: all but eight of our mutants showed AP for at least one pair of resources, and each mutant showed AP for a median of 6 resource pairs (out of 55). Finally, since our lines evolved under very weak selection, we were able to explore not only highly beneficial mutations, but the entire DFE for the occurrence of pleiotropy. This in turn allowed us to measure pleiotropic effects of a large set of mutations, making it possible to empirically test the relationship between fitness effect size and AP incidence.

We also note some important limitations of our work. First, to minimize false positive cases of pleiotropy due to error in measuring growth rates, we considered that all mutations showing <5% change from the ancestor were neutral. Effectively, we may have thus ignored mutations with effect sizes <5%, potentially underestimating the incidence and effect sizes of antagonistically pleiotropic mutations. However, this seems unlikely because we found that for many resources, small-effect mutations are depleted in AP. Second, we measured the incidence and nature of pleiotropy only for metabolic traits; specifically, for carbon utilization. Although we measured many more traits than previous studies, this is still a small fraction of traits that are probably relevant for ecological and evolutionary processes in bacteria. It is possible that antagonistic pleiotropy may be more frequent across diverse traits, such as those related to metabolism vs. stress response. Despite these limitations, our work represents the largest systematic analysis of single step mutational effects, and thus represents an important test of long-held assumptions in evolutionary biology.

In summary, we provide new insights into the incidence, nature and effect sizes of pleiotropic mutations. Although phenotypic tradeoffs clearly influence many biological processes, we suggest that at the genetic level, tradeoffs are rarer than expected. Antagonistic pleiotropy is thought to underlie the evolution and maintenance of generalists: AP may impose a cost of specialization on resource specialists, such that in heterogeneous environments, generalists that do not pay this cost are favoured (Cooper and Lenski 2000; Gompert and Messina 2016). Our results suggest that this broadly intuitive explanation needs to be more nuanced, because the incidence of AP varies significantly across environments. Thus, a generic “cost of specialization” cannot always explain the occurrence of generalists, but may have explanatory power in specific heterogeneous environments that include resource pairs showing high incidence of AP. We hope that empirical quantification of the incidence and magnitude of AP across various organisms, environments, age classes, and genetic backgrounds will provide further insights into these issues. Ultimately, we need to integrate across mechanistic and phenotypic effects to better understand the role of tradeoffs in evolution.

## ACKNOWLEDGEMENTS

We thank Olivier Tenaillon for pointing out the necessity to determine the null expectation for the incidence of pleiotropy; Santiago Elena for discussion; and members of the Agashe lab for constructive comments on the manuscript. We are grateful to Awadhesh Pandit from the Next Generation Genomics Facility at the National Centre for Biological Sciences (NCBS) for his help with whole genome sequencing, and we thank Gaurav Diwan and Aalap Mogre for their help while writing scripts for sequence analysis. We acknowledge funding and support from the National Centre for Biological Sciences (NCBS-TIFR), the Wellcome Trust/DBT India Alliance Fellowship (grant number IA/I/17/1/503091 to DA), and the Council for Scientific and Industrial Research (CSIR) of India (Senior Research Fellowship to MS).

## AUTHOR CONTRIBUTIONS

MS designed and conducted experiments, analysed data, and wrote the manuscript. JJM conducted experiments. DA conceived and designed experiments, obtained funding, analysed data, and wrote the manuscript.

